# GATA transcription factor in common bean: a comprehensive genome-wide functional characterization, identification, and abiotic stress response evaluation

**DOI:** 10.1101/2023.04.08.536124

**Authors:** Mohamed Farah Abdulla, Karam Mostafa, Abdullah Aydin, Musa Kavas

## Abstract

The GATA transcription factor has been extensively studied for its regulatory role in various biological processes in many plant species. The functional and molecular mechanism of GATA TFs in regulating tolerance to abiotic stress has not yet been studied in the common bean, a popular and commercially important crop affected by the intensifying climate change. This study analyzed the functional identity of the GATA gene family in the P. vulgaris genome under different abiotic and phytohormonal stress. The *GATA* gene family was systematically investigated in the *P. vulgaris* genome, and 31 *PvGATA* TFs were identified. The collinearity between the common bean, rice, and *Arabidopsis* was studied, and collinearity with Arabidopsis was higher than in rice. Among the 31 *PvGATAs*, 18 showed duplicated events, which suggests the importance of gene duplication in contribution to GATA gene expansion. Furthermore, all the *PvGATA* genes were classified into four major subfamilies, with eight, three, six, and 13 members in each subfamily (subfamilies I, II, III, and IV), respectively. A single GATA domain was also present in all PvGATA protein sequences, however, members of subfamily II contained additional domains like CCT and tify domains. The study also predicted and analyzed the promoter cis-regulatory elements (CREs). A total of 799 CREs was predicted in the *PvGATAs*, categorized into three categories, namely growth and development regulatory elements, stress-responsive elements, and phytohormone-responsive elements. The growth and development regulatory elements had the largest share of CREs in the promoter region of *PvGATA* genes. Additionally, we used qRT-PCR to investigate the expression profiles of five *PvGATA* genes in the common bean roots under abiotic and phytohormone treatments. The results suggest that *PvGATA1/10/25/28* may play crucial roles in regulating plant resistance against salt and drought stress at 24 hours and may be involved in phytohormone-mediated stress signaling pathways. This study provides a comprehensive analysis of the *PvGATA* gene family, which can serve as a foundation for future research on the function of *GATA* Tfs in abiotic stress tolerance in common bean plants.

## Introduction

*Phaseolus vulgaris* L., commonly known as common bean, navy, pinto, red kidney, or French beans, is a significant source of protein for global populations. The crop’s popularity stems from its noncentric and flavourful nature and adaptability (Gepts, Osborn, Rashka, & Bliss, 1986; Myers & Kmiecik, 2017). Moreover, the common bean is a vital component of healthy diets owing to its exceptional nutritional content and functional properties (Yang, Gan, Ge, Zhang, & Corke, 2018). The common bean is cultivated under different climatic conditions and produced in developed and developing countries (Beaver & Osorno, 2009). Recently, many researchers have conducted studies on common beans’ protein, fat, fatty acid, mineral content (Celmeli et al., 2018), polyphenols (Yang et al., 2018), vitamin and mineral content (AUGUSTIN, BECK, KALBFLEISH, KAGEL, & MATTHEWS, 1981), trypsin inhibitors activity (TIA), phytic acid, tannins, ascorbic acid, thiamine, and protein (Sangronis & Machado, 2007) in terms of food security and nutrition. One of the most critical challenges facing the world in the next three decades is achieving food security. Moreover, climate change is likely to have an adverse impact on agricultural production and primary crop nutrition. Extensive crop modeling studies have demonstrated that common bean growing regions and yields will be negatively affected by 2050 (Hummel et al., 2018).

The growing number of next-generation sequencing (NGS) platforms has made numerous plant genomes available, presenting an opportunity to understand the evolutionary history of plants and investigate the role of transcription factors (TFs) in plant biology. TFs are proteins that control the transcription rate by binding to cis-acting elements in specific gene promoter regions, playing essential roles in plant developmental processes, hormone signaling networks, and disease resistance responses. Plant genomes contain various transcription factor families, including WRKY, MYB, DREB, bZIP, MADS-box, bHLH, and GATA. GATA TFs are highly conserved among eukaryotic organisms and are essential in plant development and response to environmental stresses. These TFs have a DNA-binding domain that recognizes the consensus motif WGATAR, present in the promoter regions of their target genes. Recent studies have shown that GATA genes play crucial regulatory roles in several biological processes, such as seed germination, embryogenesis, chloroplast development, flowering time, carbon and nitrogen assimilation, and response to abiotic and biotic stresses. In this regard, GATAs participate in plant developmental processes and respond to environmental challenges by binding to DNA regulatory areas to control their downstream genes.

The availability of numerous plant genomes due to the rising number of next-generation sequencing (NGS) platforms has created an opportunity to comprehend the evolutionary history of plants and determine the role of transcription factors (TFs) in plant biology. TFs are proteins that control the transcription rate by binding to cis-acting elements in specific gene promoter regions, playing essential roles in plant developmental processes, hormone signaling networks, and disease resistance responses (Kim, Xi, & Park, 2021). Plant genomes contain various transcription factor families, including WRKY (J. Jiang et al., 2017), MYB (Ambawat, Sharma, Yadav, & Yadav, 2013; Kavas et al., 2022), DREB (Khan, 2011), bZIP (Alves et al., 2013), MADS-box (de Folter et al., 2005), bHLH (Kavas et al., 2016; Liang et al., 2023), and GATA (Kim et al., 2021; Reyes, Muro-Pastor, & Florencio, 2004; Wang et al., 2023). GATA transcription factors (TFs) are evolutionarily conserved transcription and are ubiquitous among eukaryotic organisms such as mammals, fungi, and plants (Dobrzycki, Lalwani, Telfer, Monteiro, & Patient, 2020; J. A. Lowry & W. R. Atchley, 2000; Romano & Miccio, 2020).

GATA transcription factors are known to play essential roles in plant development and response to environmental stresses. These transcription factors are characterized by a highly DNA-binding domain that recognizes a consensus motif WGATAR (W = T or A; R = G or A) in the promoter regions of their target genes (J. A. Lowry & W. R. Atchley, 2000; Reyes et al., 2004). Recent studies have shown that GATA genes play crucial regulatory roles in several biological processes (Schwechheimer, Schröder, & Blaby-Haas, 2022)., such as seed germination, embryogenesis, chloroplast development, flowering time, carbon and nitrogen assimilation, and response to abiotic and biotic stresses (An et al., 2019; C. Jiang et al., 2021; L. Jiang, Yu, Chen, Feng, & Li, 2020; H. Liu et al., 2019). In this regard, GATAs participate in plant developmental processes and respond to environmental challenges by binding to DNA regulatory areas and controlling genes downstream of the pathway (C. Jiang et al., 2021).

Furthermore, The *GATA* TFs have a significant function in diverse abiotic stresses in plants. Numerous *GATA* factors were found to be up-regulated or downregulated in soybean leaf in response to low nitrogen stress. Two *GATA* factors, *GATA44* and *GATA58*, were shown to be involved in nitrogen metabolism regulation in soybean (C. Zhang et al., 2015). Under salinity conditions, *OsGATA8* regulates the expression of critical genes involved in stress tolerance, scavenging reactive oxygen species, and chlorophyll biosynthesis (Nutan, Singla-Pareek, & Pareek, 2020). Under different nitrogen levels, the poplar *GATA* transcription factor *PdGNC* can control chloroplast ultrastructure, photosynthesis, and vegetative development in Arabidopsis (An, Han, Tang, Xia, & Yin, 2014). Therefore, *GATA* transcription factors are important regulators of plant growth and development, as well as adaptation to different environmental conditions.

Because of little information on the *GATA* genes and their role under abiotic stress in *Phaseolus vulgaris* is available. Herein, we performed genome-wide identification, phylogenetic studies, expression level analysis, and identification of amino acid patterns and transmembrane helix of 32 *GATA* genes in common bean. These findings enrich our knowledge about *PvGATA* genes.

## Material and methods

### Retrieval and characterization *of GATAs* in *P.vulgaris*

A comprehensive and non-redundant data set of common bean proteins containing the conserved GATA domain was compiled using BLAST and keyword searches. Firstly, The Hidden Markov Model (HMM) and BLASTP algorithms were used to identify pvGATA proteins (Finn et al., 2015). Additionally, we conducted a keyword search in the Phytozome v13 database (https://phytozome-next.jgi.doe.gov/) to identify additional possible GATA genes that were missed in the first step. The candidate sequences of GATAs were confirmed using InterPro (https://www.ebi.ac.uk/interpro/) and Pfam (http://pfam-legacy.xfam.org/) tools (Hunter et al., 2008; Mistry et al., 2020). The query proteins and nucleotide sequences of all putative *pvGATA* genes were obtained from (https://phytozome-next.jgi.doe.gov/). Putative *pvGATA* genes were named based on their arrangement order on chromosomes of the *P.vulgaris* genome. Moreover, the length of amino acids, molecular weights (MW), and isoelectric point (pI) of GATA proteins were calculated using ExPASy (http://www.expasy.ch/tools/pi_tool.html). The PROSOII tool was used to predict the solubility of candidate GATA proteins based on their sequences (Hon et al., 2021).

### Multiple sequence alignment and phylogenetic analysis

The protein sequence of each putative GATA genes from *Phaseolus vulgaris, Arabidopsis thaliana, and Oryza sativa* were acquired from the Phytozome v13 database for the Phylogenetic structural analysis and synteny analysis inter and intra-species. A phylogenetic tree derived from multiple sequence alignment was carried out by the ClustalW 2.0 program in MEGAX software. The neighbor-joining method-based phylogenetic tree was constructed using the bootstrap test (1000 replicates) and the Jones-Taylor-Thornton (JTT) model in the IQtree web tool. Phylogenetic trees were visualized with ITOL v3 (http://itol.embl.de/).

### Chromosomal localization, Motifs, CREs, and gene structures in common bean

Using a generic feature format version 3 (GFF3) obtained from the Phytozome v13 database, the chromosomal position of *PvGATA* genes was examined. The TBTools software was employed to locate *PvGATA* genes on their corresponding chromosome locations. We examined the gene structure (exons and introns) as well as conserved domains and motifs to demonstrate the potential link between the evolution process of *PvGATA* genes and their structure-function correlation. TBtools (T. Chen et al., 2011) was utilized to visualize the motifs of *Phaseolus vulgaris* GATA proteins predicted and analyzed by MEME. The selected parameters specified that each sequence could contain zero or one contributing motif site, with a total of six repeated motifs chosen. The widths of the motifs were set to a range of 6 to 50, while the remaining parameters were kept at their default values (Wu et al., 2016). To ensure accuracy, each motif was individually evaluated, and only those with an e-value of less than 1e-10 were considered for motif detection in *Phaseolus vulgaris* GATA proteins. To analyze the cis-acting regulatory elements, the upstream 2000 DNA sequences from the translation initiation site of each *PvGATA* gene were extracted from Phytozome v13 database and submited to the PlantCare web tool (http://bioinformatics.psb.ugent.be/plantcare/html). The results were then visualized using the tbtools software.

### Analysis of the evolutionary divergence of PvGATA genes family using Gene Duplication, Synteny Analysis, and Protein-protein interaction

In this study, we utilized the One Step MCScanX method to perform multiple alignments using the genome sequence file and the genome structure annotation file of *P. vulgaris* downloaded from Phytozome v13 database. We used the dual synteny plotter for the MCScanX program to visualize the collinearity results. These steps were conducted using the tbtools software (C. Chen et al., 2020). To predict gene duplications of the *PvGATAs*, we employed the Plant Duplicate Gene Database (PlantDGD) and identified tandem repeats among duplicate genes located on the same chromosome (Qiao et al., 2019). Additionally, we conducted a BLASTP search to detect segmental duplications of the PvGATA proteins in the common bean and used the MCScan tool to determine their collinear blocks. To estimate the evolution of the *PvGATA* genes, we calculated the ratio between nonsynonymous mutation rate and synonymous mutation rate (Ka/Ks) via TBtools software based on previous reports (Kavas et al., 2022). In short, we first blasted PvGATA proteins against the Phytozome v13 database and filtered the hits over 60% sequence similarity threshold to estimate the evolutionary relationship of these proteins. We then created a tab-delimited text file to calculate Ka/Ks in TBtools software. To analyze the protein-protein interaction (PPI), we utilized the STRING STRING web portal (http://string-db.org), and orthologs of these proteins were found in *Arabidopsis thaliana* (Szklarczyk et al., 2018). Finally, we investigated the PPI interaction of these proteins based on default settings.

### Plant material and qRT-PCR transcript expression analysis

This study used a locally cultivated commercial common bean variety (*Ispir*) for its known resistance to saline environments. *Ispir* seeds were surface sterilized in a 5% sodium hypochlorite solution before being planted in vermiculite-filled pots. Plants were cultivated in a fully controlled growth chamber at 24 C with a photoperiod of 16 hours light and 8 hours dark. After four weeks, the plants were treated to drought, salinity, and phytohormones (Indole acetic acid and Abscisic acid) treatments using polyethylene glycol 6000 (PEG), NaCl, ABA, and IAA, respectively. The plants were subjected to salt and drought stress by adding 200 mM NaCl and PEG (20%) to the Hoagland solution. Phytohormone treatment was done by spraying 100 μM ABA and 100 μM IAA on the leaves. Samples of stress and hormone-treated roots were taken at 6, 24, and 48th hours following stress treatment and preserved at -80 C until use in RNA isolation. We extracted total RNA using the RNeasy Plant Mini Kit (Qiagen) according to the manufacturer’s instructions. The purity and concentration of the RNA were verified using a NanoDrop TM 2000/2000c spectrophotometer and a 1.5% (w/v) agarose gel. The iScript™ cDNA Synthesis Kit (Bio-rad, USA) was used to create the first strand cDNA. Phytohormone and stress-related expression of *five randomly selected candidate PvGATA* genes were measured with qRT-PCR analysis performed on the Agilent Mx3000P device with Solis BioDyne 5 × HOT FIREPol^®^ EvaGreen^®^ qPCR Mix Plus (ROX). All primers used for the qRT-PCR expression analysis are listed in Table S1. qRT-PCR conditions were carried out at 95 °C for 2 min, at 95 °C for 15 s, and 60 °C for 1 min. The 2^-ΔΔCT^ technique was used to compute the relative expression.

### RNA-Seq-based in silico gene expression analysis of P. vulgaris

In this study, the expression levels of *PvGATA* TFs were investigated using data of RNA-seq from different publicly available datasets, which had been collected from various common bean genotypes under various tissues, growth stages, and abiotic and biotic stress. Raw RNA-seq values were obtained from the Sequence Read Archive (SRA) in the NCBI database, and transcriptome analyses were conducted using cloud bioinformatic tools such as CyVerse and Galaxy. HISAT2 was used to map the reads to the reference genome, while Stringtie 1.3.3 and Ballgown were utilized for transcript assembly and differential expression analysis, respectively. DEGs were identified as genes with a fold change value log2 > 1 and a p-value < 0.05. A heatmap was generated using TBTools to visualize the log2 fold change values. For stress expression analysis, seven different comparisons were conducted to determine the expression levels of all the *PvGATA* genes in diverse tissues and under various stress conditions. Firstly, two comparisons were made using RNA-seq data (PRJNA656794) from leaf, and root explants of resistant (Ispir) and sensitive (T43) genotypes treated with salt stress. The third comparison was conducted using data from salt-stressed lower hypocotyl at the sprout stage of the Ispir genotype (PRJNA691982). The fourth comparison was made using RNA-seq data from drought-tolerant genotype Perola under drought stress versus control (PRJNA508605), while the fifth comparison was conducted using RNA-seq read values obtained from *P. vulgaris* plants infected with *Sclerotinia sclerotiorum* (PRJNA574280). Finally, the seventh comparison was made using the data collected from common bean plants under cold stress treatment (PRJNA793687). The tissue-specific expression patterns of PvGATAs were obtained from the publicly available transcriptome data (PRJNA210619).

## Results

### Identification and characterization of PvGATA genes

We identified and extracted 31 *GATA* transcription factor (TF) genes through a BLAST-P search of the common bean genome database in Phytozome V13 and NCBI. We removed genes that did not contain a typical GATA domain. A list of the identified genes is shown in Table 1. We renamed these GATA TFs as *PvGATA* and assigned them a numerical designation based on their order on the chromosomes. The genes encoding the 31 *PvGATA* TFs had amino acid lengths ranging from 107 (*PvGATA3*) to 544 (*PvGATA11*), with an average size of 290 amino acids. The basic physiochemical properties analysis revealed an isoelectric point (pI) ranging from 9.95 (PvGATA27) to 4.96 (PvGATA23), with an average of 7.28 and an average molecular weight of 31.94 kDA. The highest molecular weight was 59.84 kDA in PvGATA11, and the lowest was 11.77 kDA in PvGATA3. The Grand Average of Hydropathy (GRAVY) of PvGATA proteins was negative, ranging from -0.9 to -0.38, indicating a non-polar and hydrophilic nature. Lastly, the subcellular localization of PvGATA proteins was predicted to be in the nucleus, except for PvGATA3, which was predicted to be extracellular.

**Table 1.**
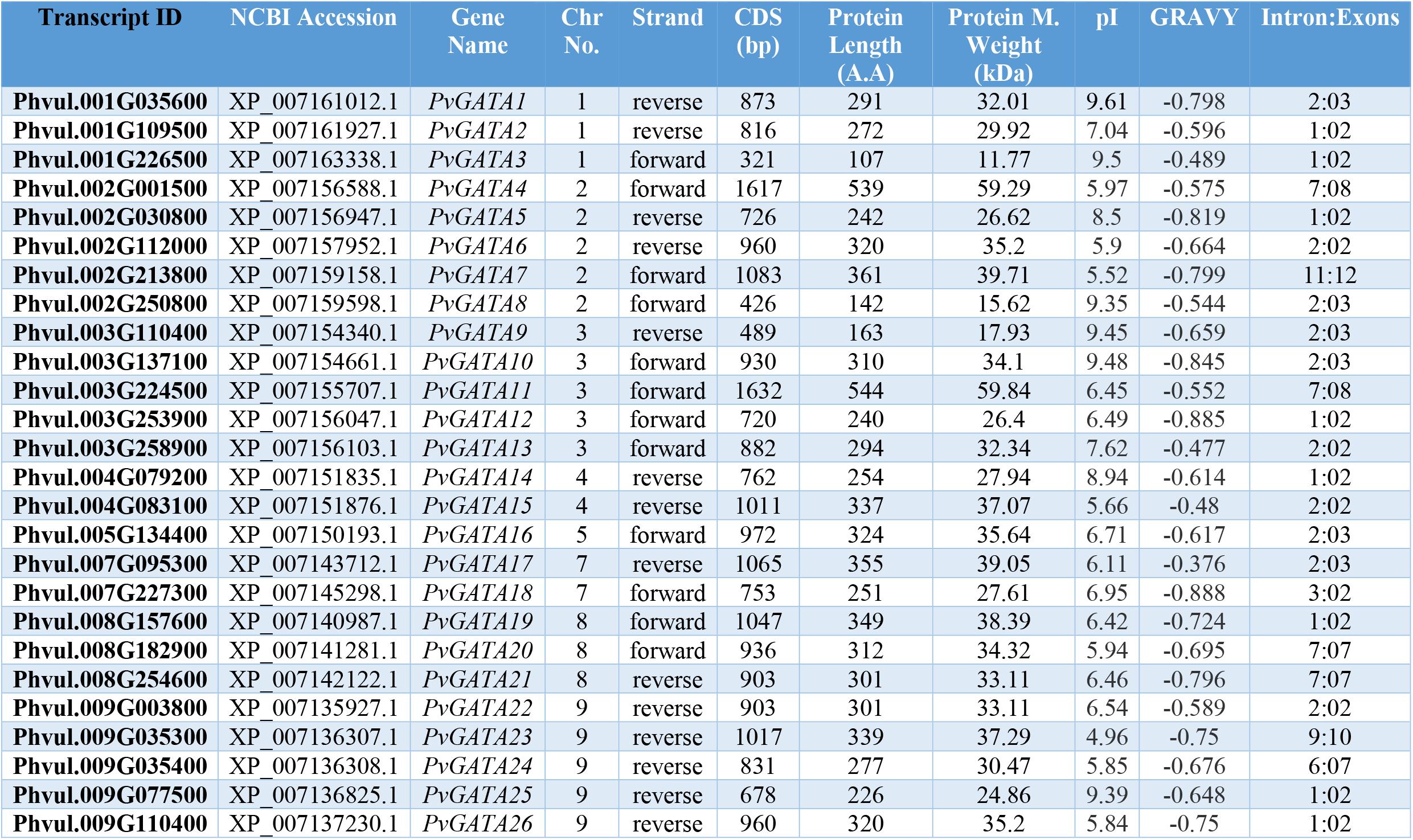
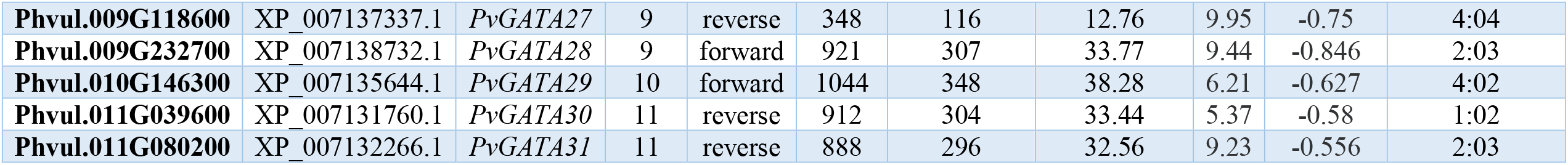
Characterization of GATA proteins

### Phylogenetic Classification of PvGATA, their motif, gene structure, and conserved domain analysis

A maximum likelihood analysis was conducted to examine the inter- and intraspecific phylogenetic relationships among GATA TF protein sequences in the common bean, *Arabidopsis*, and rice genomes. Following previous studies, the 31 PvGATA genes were grouped based on their conserved domains and motif structures (Figure 1). As anticipated, the *PvGATA* genes were divided into four sub-families, namely subfamily I-IV. The analysis revealed that 8 *PvGATA* genes were in subfamily I, three genes in subfamily II, 6 in subfamily III, and 14 in subfamily IV.

**Figure 1.**
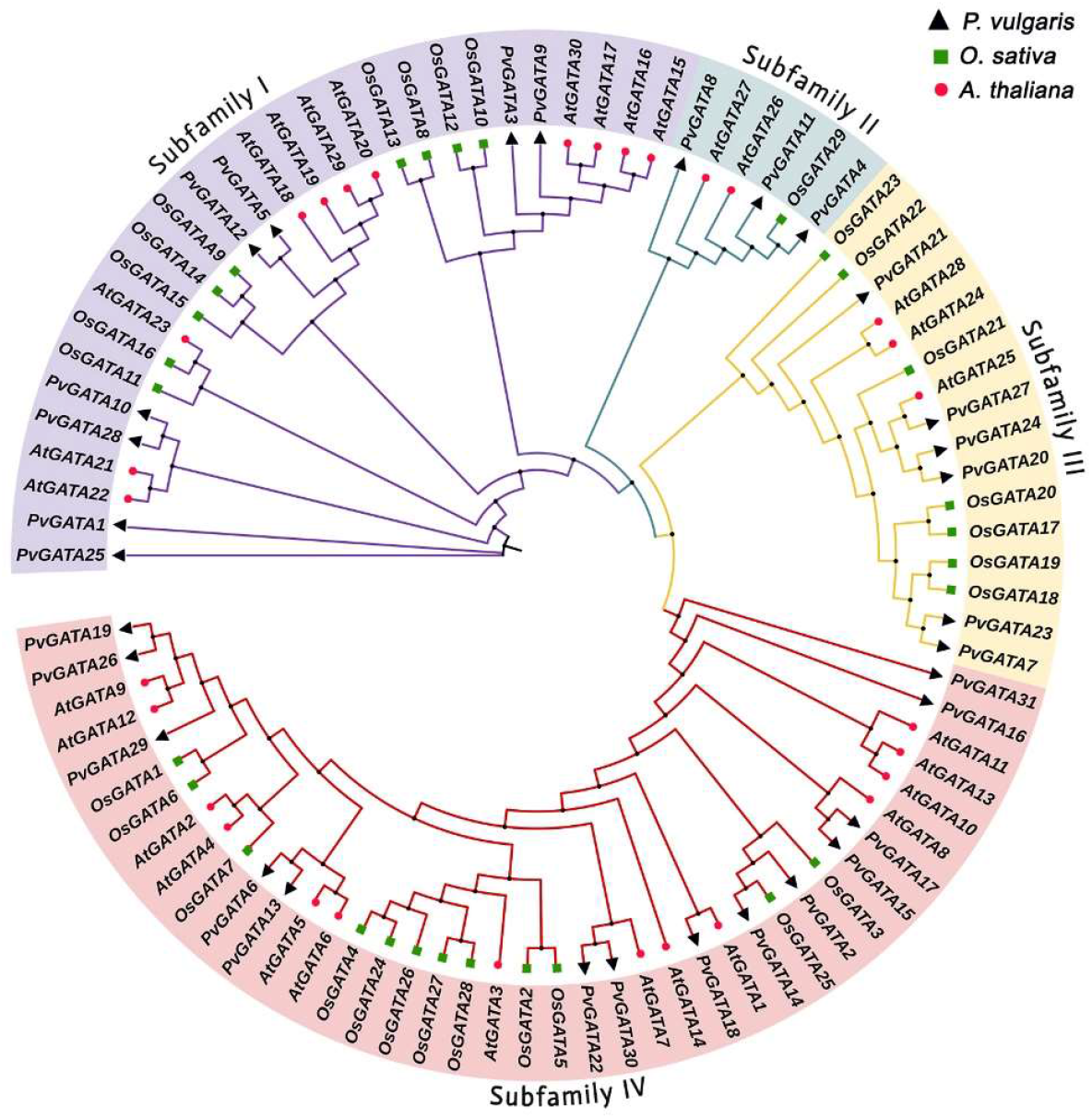
Phylogenetic relationship analysis of the GATA proteins between Phaseolus vulgaris, Oryza sativa, and Arabidopsis thaliana. the whole protein sequences of GATA gene family were used for alignment using the MEGA X software. The phylogenetic tree was constructed using IQ-TREE 2 web tool using maximum likelihood with a 1000 bootstrap replicates. Different colored branches correspond to distinct GATA subfamilies, and the GATA IDs of Arabidopsis thaliana and Oryza sativa were assigned based on previous studies.

To more clearly compare the *PvGATA* transcription factors at both the nucleic acid and protein levels, we conducted a comprehensive analysis of the conserved motifs, gene structure, domains, and the phylogenetic relationships among the 31 PvGATA proteins (Figure 2a). Using the MEME search tool, we identified six conserved motifs labeled as motifs 1-5. The results showed varying compositions and distributions of these motifs among the PvGATA proteins, with the exception of PvGATA17, which had no assigned motifs. In contrast, motifs 1 and 5 were found in all the PvGATA proteins (excluding PvGATA17) and in the same arrangement. We also used the NCBI CD blast tool to analyze the conserved domains of the PvGATA proteins and found that the GATA domain was present in all 31 proteins. The *TIFY* domain and *CCT* (*CO, COL*, and *TOC1*) domains were present in subfamily II, while the *ASXH* domain was present in subfamily IV. Our comparison of the genetic exon and intron architecture of the *GATA* genes revealed that they had a varying number of exons, ranging from 2-13. Notably, subfamilies III and IV had a higher number of introns compared to the rest of the subgroups, which had 1-3 introns. Our analysis confirmed that there were similarities in motif structure, conserved domains, and exon/intron configuration among members of the same subfamily, further supporting our phylogenetic analysis and subfamily classification.

**Figure 2.**
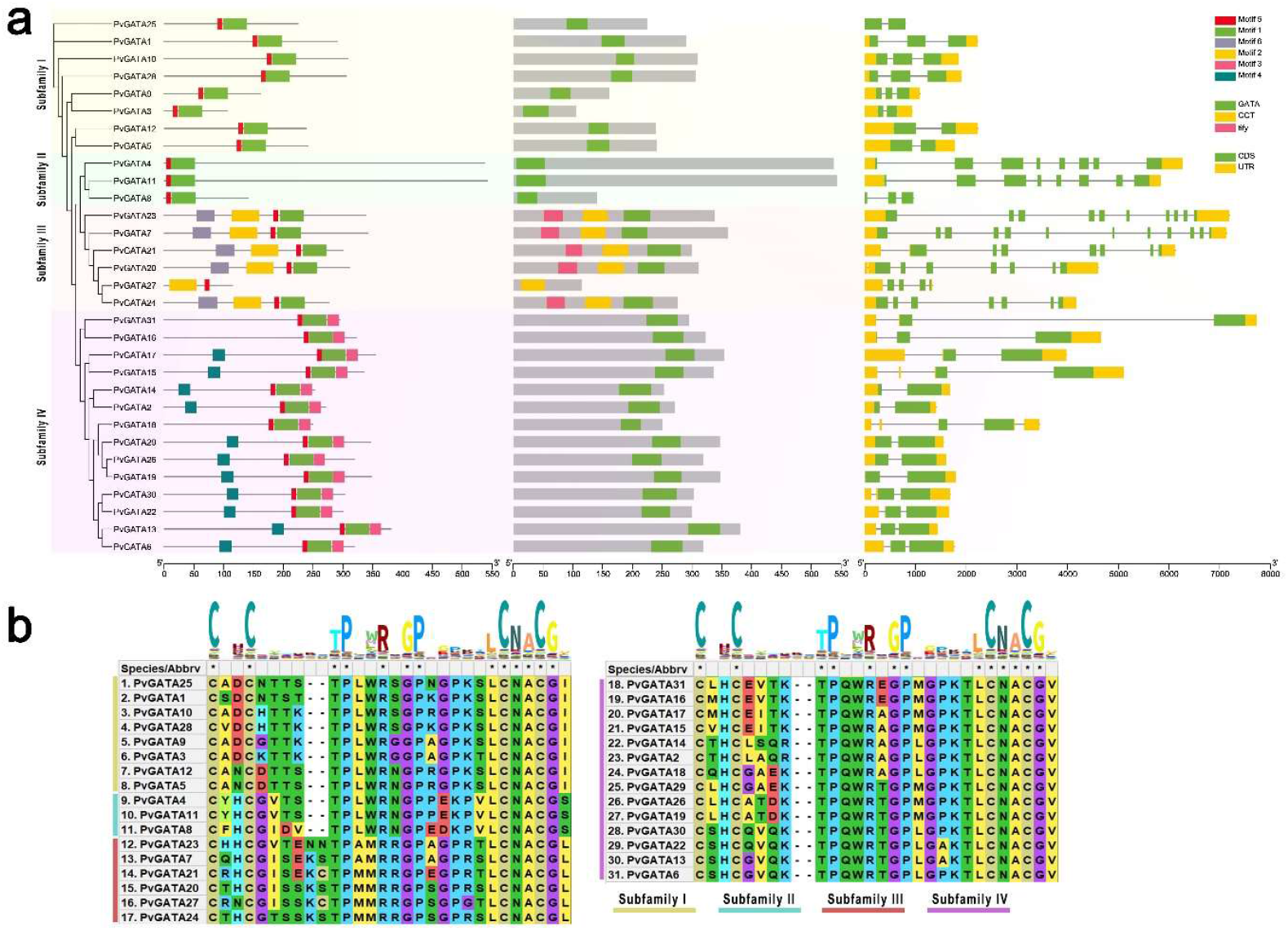
(a) The evolutionary relationships, conserved motifs, domain arrangement, and gene structures of the PvGATA TFs. A maximum likelihood phylogenetic tree was generated based on the full-length sequences of the PvGATA, and 1000 bootstrap replicates using the IQTree webtool. The distribution of conserved motifs in PvGATA was predicted and was limited to 6 conserved motifs. Three conserved domains were found by analyzing the conserved domain structure of PvGATA sequences using the NCBI CD database. Moreover, the gene structures of PvGATA were analyzed and visualized, including the introns (black lines), exons (CDS, green rectangles), and untranslated regions (UTRs, yellow rectangles). The scale bar represented 100 bp. (b) The distribution and conserved regions of the GATA domain were investigated across all PvGATA proteins with respect to subfamilies using MEGA X software, with the HHM logo of the GATA domain shown.

Furthermore, the GATA domain analysis yielded similar results to those found in *Arabidopsis* and rice (Reyes et al., 2004). All the subfamilies I-IV of PvGATAs exhibited 13 conserved residues in the zinc finger loop (C-X2-C-X18-C-X2-C). All 8 PvGATAs in subfamily I contained 21 residues along the residues in the zinc finger (C-X2-C-X20-C-X2-C) (Figure 2b). Subfamily II had 22 conserved residues, subfamily III had 21 conserved residues, and subfamily IV contained 22 conserved residues. Moreover, several amino acid sites within the GATA domains, such as LCNACG residues, demonstrated high levels of conservation among common bean, *Arabidopsis*, and rice.

### Chromosomal localization, synteny analysis, and PPI of *PvGATA* gene family

To understand the chromosomal distribution of the common bean *GATA* genes, we physically mapped the locations of the 31 *PvGATA* genes on the common bean genome (Figure 3a). The distribution was uneven, with Chr09 having the highest number of *PvGATA* genes (7), followed by Chr02 and Chr03, each containing 5 *GATA* genes. *PvGATA* genes were located on Chr01 and Chr08, with three genes on each chromosome. Chr04, Chr07, and Chr11 each contained 2 *PvGATA* genes, while the minimum number of 1 *PvGATA* gene was found on Chr05 and Chr10.

**Figure 3.**
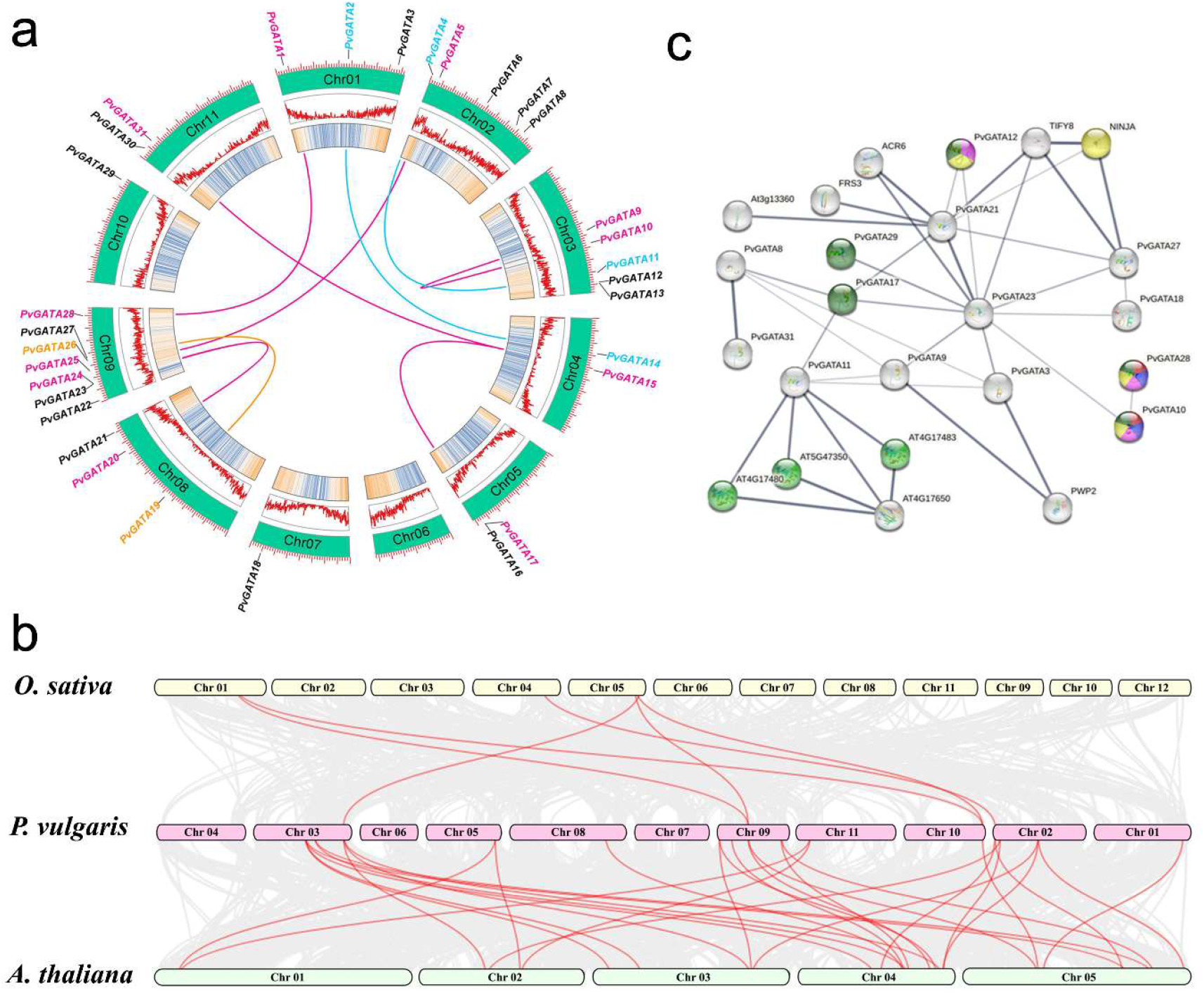
a: Chromosomal localization and synteny analysis of PvGATA proteins in the common bean genome. Genes IDs in black indicate the absence of collinearity, genes, and lines colored in magenta indicate dispersed duplication; cyan indicates whole genome duplication, and golden colored lines indicate transposed duplicated pairs. The two rings in the center represent the chromosome’s gene density. b. Collinearity relationship analysis between P. vulgaris to O. sativa and A. thaliana. Gray lines indicate all synteny blocks found between the genomes of the species, and red lines indicate the gene pairs with duplicated events. c. Protein-protein interaction analyses were performed on the String web tool. The thickness of the lines represents the reliability of the results. The various functional associations are represented by different colors, including yellow for the regulation of shoot development, red for the negative regulation of gibberellic acid-mediated signaling pathway, blue for the negative regulation of seed germination, purple for the regulation of flower development, and green for positive regulation of nitrogen compound metabolic process.

The present study employed the MCScanX method to investigate the *PvGATA* gene duplication events. Specifically, we observed nine gene pairs exhibiting duplication events. Six gene pairs were recorded to have dispersed duplication, two gene pairs had whole genome duplication, and one gene pair was identified to have undergone transposed gene duplication event. To explore the evolutionary process and selection pressure of the *PvGATA* genes, we calculated the substitution rate ratio Ka/Ks for all the 31 GATA genes present in the common bean genome, as shown in (Table 2). Our results indicate that the Ka/Ks ratios for all the gene pairs were less than one and ranged from 0.381 to 0.126, with the gene pairs *PvGATA4*-*PvGATA11* and *PvGATA1*-*PvGATA28* exhibiting the highest and lowest ratios, respectively. These findings suggest that the *PvGATA* gene pairs may have undergone purifying selection during evolution and played a crucial role in maintaining the conserved structure of the *PvGATA* gene.

**Table 2.**
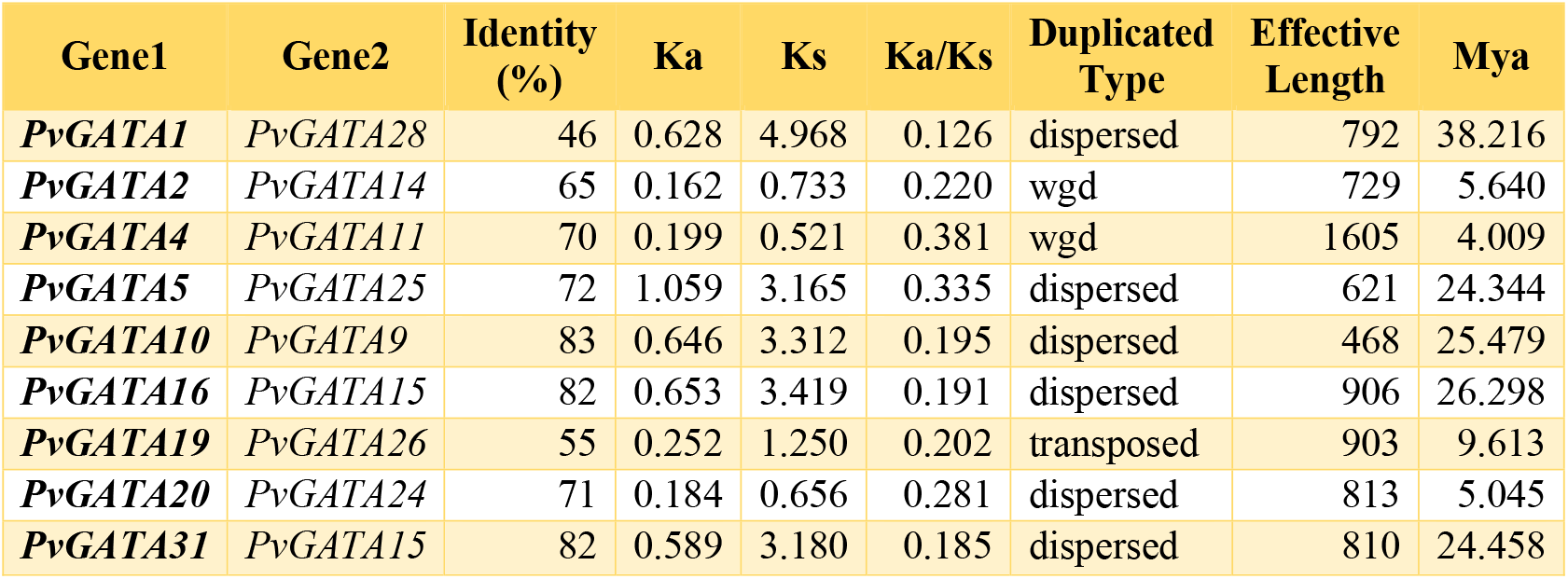
Inter-specific gene duplication analysis of PvGATAs

Based on protein-protein interaction (PPI) analyses, several GATA proteins, including PvGATA10, PvGATA12, PvGATA17, PvGATA28, and PvGATA29, have been found to play positive roles in the nitrogen compound metabolic process. On the other hand, PvGATA21 interacts with TIFY8 and NINJA, which are negative regulators of jasmonic acid signaling. PvGATA10 and PvGATA28 have been observed to have a negative role in the gibberellic acid-mediated signaling pathway and seed germination. Furthermore, PvGATA10, PvGATA12, and PvGATA28 are involved in shoot system development and flower development. These findings shed light on the diverse functions of PvGATA proteins and their potential roles in various physiological processes in common bean plants.

### Cis-Regulatory elements in the promoter region of *PvGATA*

To predict and analyze the promoter *cis*-regulatory elements (CREs) in *PvGATAs*, we extracted the 2000 bp nucleotide sequences upstream of the genes and submitted them to the PlantCARE database. A total of 799 CREs were predicted in all *PvGATAs*, with *PvGATA12* having the most (51) and *PvGATA14* having the least (12) number of CREs. These *cis*-Regulatory elements were categorized into three categories, namely Growth and Development regulatory elements, Stress-responsive elements, and Phytohormone-responsive elements (Figure 4b, Table S2).

**Figure 4.**
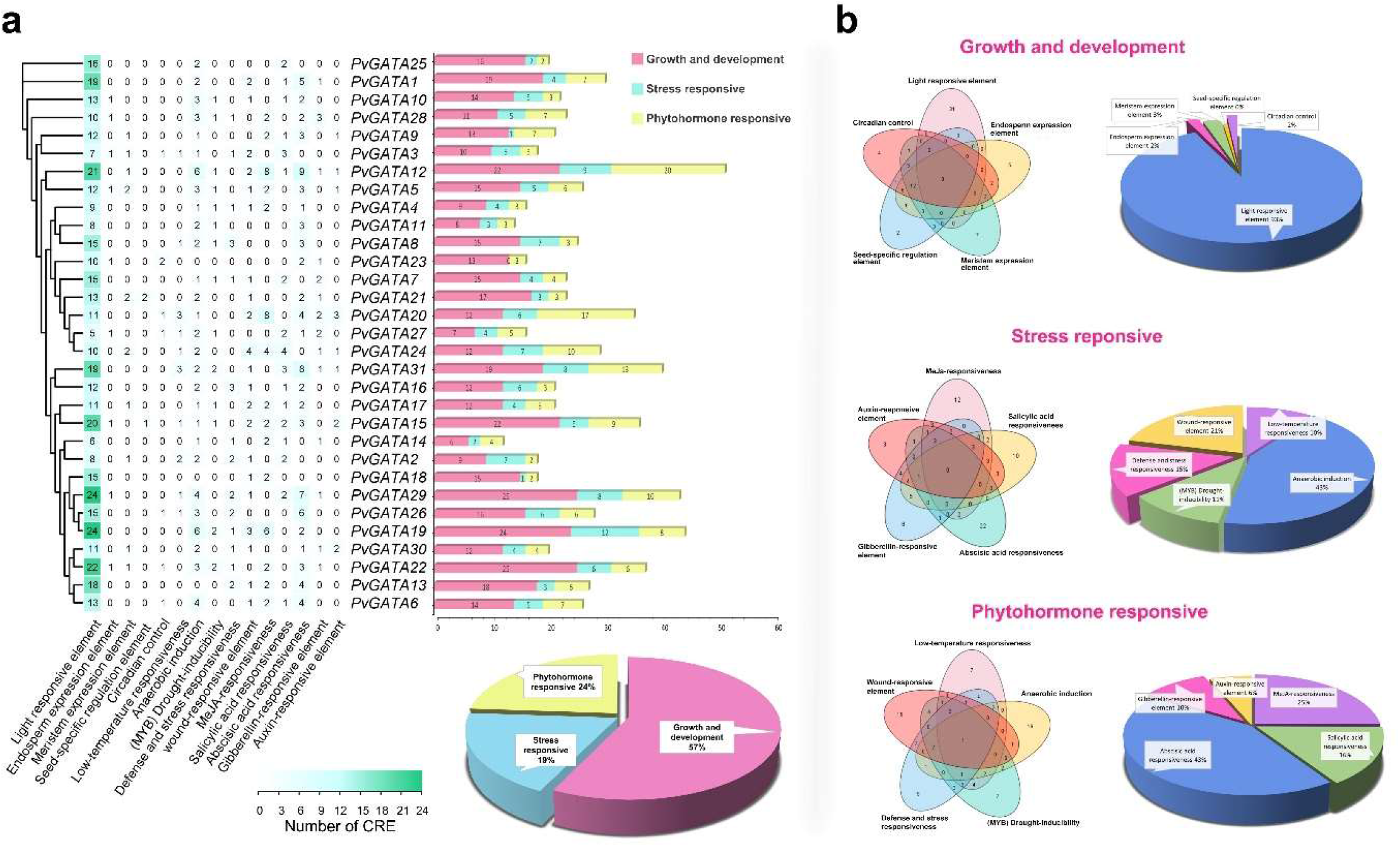
Analysis of cis-regulatory elements (CREs) in the putative promoter region of PvGATA genes using the PlantCARE database. (a) The number of predicted CREs located in the 2000 bp upstream of the PvGATA genes and the distribution of the three categories of CREs among the members of the PvGATA gene family. (b) Venn diagram and a pie chart showing the distribution of different functional categories of CREs identified in the PvGATA promoter region.

i. Growth and development regulatory elements had the largest share of CRE in the promoter region of *PvGATA*, with a total of 457 elements and 57.20% of total CREs. All the subfamilies in *PvGATA* genes carried elements from this category except subfamily II, which only had elements associated with light responsiveness. Within this category, CREs light responsiveness (Box 4, LAMP element, GATA-motif, ATCT-motif, etc.) was heavily abundant, totaling 424 elements and 93% of the category’s total elements. Other elements in this category included meristem expression elements (CAT-box) with 13 elements (3%), endosperm expression (GCN4_motif) with nine elements (2%), circadian control with eight elements (2%) present in all PvGATA subfamilies except subfamily II and III, and seed-specific elements with three elements present in the promoters of *PvGATA15* and *PvGATA21* only.
ii. The second category is stress-responsive CREs. A total of 151 (18.9%) CREs were predicted in the promoter region of *PvGATAs*. At least one element was present in all the *PvGATA* TFs except in *PvGATA23*. The stress-responsive category includes anaerobic induction elements (ARE), which had the most elements, with 65 (43%) predicted, followed by wound-responsive elements (WRE) with 32 elements (21%), defense and stress responsiveness with 22 elements (15%), (MYB) drought inducibility elements with 17 elements (11%), and low-temperature responsiveness (LTR) with the least number of elements, only 15 (10%).
iii. The third category of CREs identified in the promoter region of *PvGATAs* is related to phytohormone response, with a total of 191 (23.9%) predicted CREs. The most common phytohormone-responsive element is the abscisic acid responsiveness element, with 82 (43%) CREs, followed by methyl jasmonate (MeJA)-responsive elements with 48 (25%) elements, salicylic acid-responsive elements with 30 (16%) CREs, gibberellin-responsive elements with 19 (10%) CREs, and auxin-responsive elements with the least number of CREs, only 12 (6%). Interestingly, subfamily II lacks CREs associated with gibberellin, auxin, and salicylic acid-responsive elements, suggesting the absence of these genes in the hormone regulation network.

### Functional GO annotation of *PvGATA* TFs

The functional annotation of *PvGATA* transcription factors was analyzed using the Blast2Go plugin in the OmicsBox software. The analysis revealed 17 gene ontology annotations in all *PvGATA* TFs, which were categorized into three ontologies: molecular function, biological process, and cellular component (Figure 5, Table S3). The molecular functions of *PvGATAs* were associated with protein, ion, metal, and DNA binding activities, which is consistent with the known association of *GATA* TFs with DNA binding activities. The cellular components of *PvGATAs* were predominantly located in the intercellular space, membrane, and nucleus, which underscores their importance in the development of common bean plants. Under biological process annotations, all *PvGATAs* were potentially involved in regulating metabolic processes such as nitrogen, organic compounds, and primary metabolites. Some *PvGATA* TFs were also associated with the positive regulation of cellular processes, as well as the development and regulation of cellular processes. Overall, these findings highlight on the potential functions of *PvGATAs* in the development of common bean plants.

**Figure 5.**
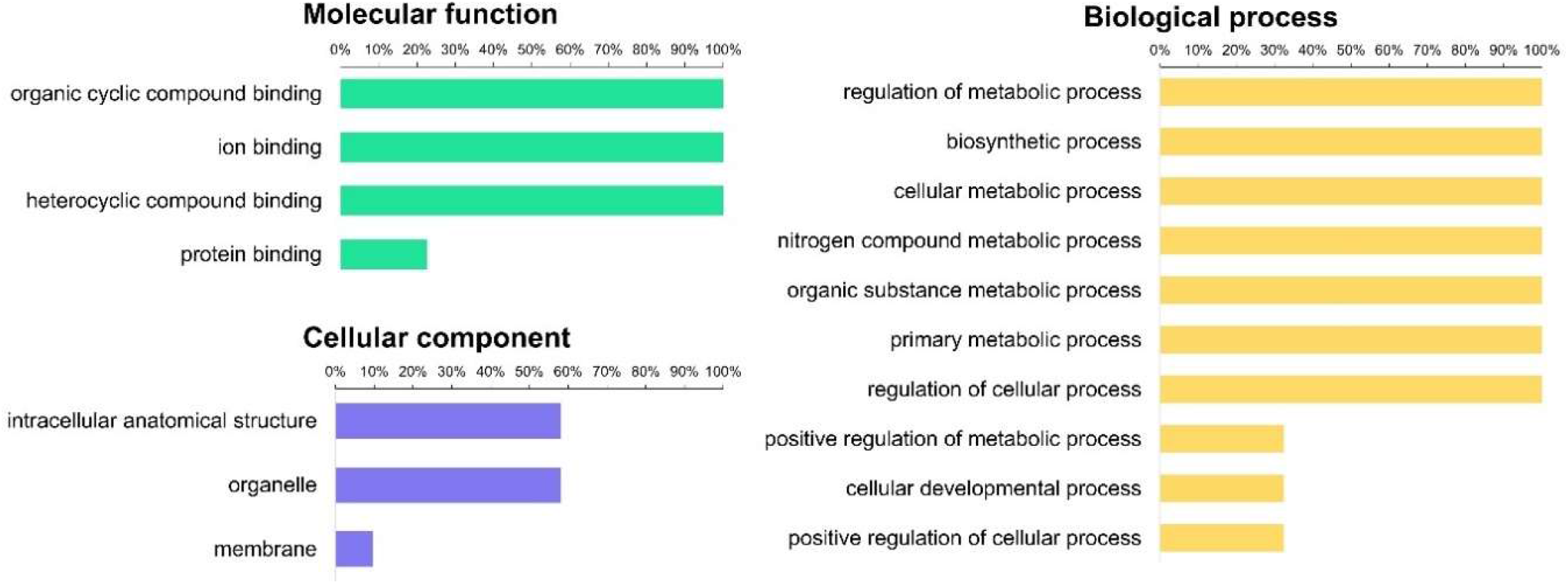
Gene ontology (GO) analysis of the PvGATA gene family using Blast2GO software. The distribution of GO annotations was determined for the genome-wide PvGATA gene family. The bar graph shows the percentage of PvGATA sequences assigned to different biological processes based on GO annotations, including molecular function, cellular component, and biological process.

### Expression analysis of *PvGATA* TFs on subcellular levels, organ, and stress conditions

In order to investigate the subcellular localization of the PvGATA proteins, we utilized the WoLF PSORT online database. Our results revealed that all PvGATA proteins, except PvGATA3 and PvGATA8, were predominantly localized in the nucleus (Figure 6a, Table S4). Precisely, members of subfamily IV, such as PvGATA2, PvGATA6, and PvGATA15, and subfamily III, such as PvGATA23 and PvGATA7, were predicted to be 100% localized in the nucleus. Furthermore, a subset of PvGATA proteins was predicted to localize to other subcellular organelles. For instance, members of subfamily II, including PvGATA4 and PvGATA11, were found to be significantly present in the chloroplast, while PvGATA8 was predicted to be localized in the cytoplasm and extracellular organelles.

**Figure 6.**
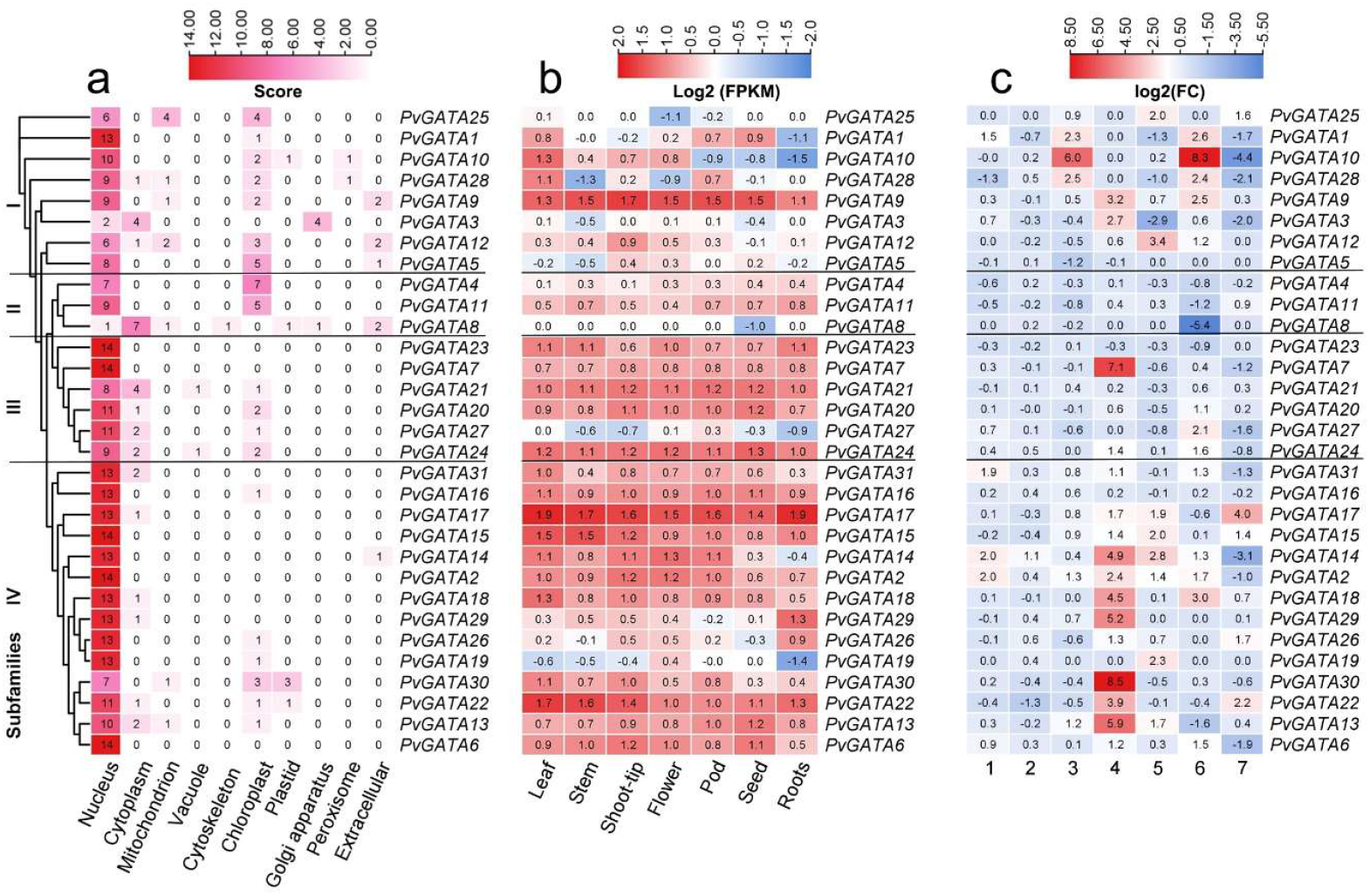
Heatmaps were generated to examine the expression patterns of PvGATA TFs under various cellular compartments, developmental stages, and stress conditions. The heatmaps were constructed and visualized using TBTools software. Black lines are used to differentiate between the subfamilies. (a) the sub-cellular localization of PvGATA proteins was predicted using the WoLF PSORT web tool. (b) The tissue-specific expression profiles of PvGATA at different developmental stages of the common bean plant were analyzed using publicly available transcriptome data (PRJNA210619) and displayed in a heatmap. The normalized fragments per kilobase of transcript per million fragments (FPKM) values were transformed by Log2(FPKM). (c) Expression analysis of PvGATA transcripts on different biotic and abiotic stress conditions using RNA-seq data publicly available. 1 and 2 samples were collected from the leaves and roots of a salt-tolerant vs. salt-sensitive cultivar under salt-stress conditions. (PRJNA656794). 3, Samples from lower hypocotyl under salt stress after 12h vs. control (PRJNA691982). 4 drought stress of resistant cultivar vs. control at 150mins after treatment. 5, Sensitive cultivar under drought stress vs. control (PRJNA508605). 6, fungal plant pathogen infected vs. non-infected samples of the resistant cultivar (PRJNA574280). 7, low-temperature stress of resistant common bean plant vs. control treatment (PRJNA793687).

We investigated the expression profile of GATA transcription factors (TFs) in various tissues of the common bean plant using publicly available transcriptome data (O’Rourke et al., 2014). The tissues analyzed included leaf, stem, shoot tip, apical meristem, young flower, young flower pods, seeds, and whole roots with root tips. As shown in Figure 6, we observed significant differences in the expression patterns of *PvGATA* transcripts among the subfamilies (Table S4). In subfamily I, almost all *PvGATA* genes, except for *PvGATA5*, were highly expressed in vegetative leaf tissues. In contrast, *PvGATA9* was highly expressed in all tissues analyzed, with the highest expression in shoot tips, suggesting its crucial role in tissue development. The expression profile of subfamily I members was considerably downregulated or had no expression in root and root-tip tissues, except for *PvGATA9*. In subfamily II, gene expression was moderate and less than 0.8, indicating their subordinate role in organ development. In subfamily III, a uniform preferential expression pattern was noted from all members, except for *PvGATA27*, which was negatively regulated on roots, stem, and shoot tips. Gene members in subfamily IV were preferentially expressed in vegetative and flower tissues. Among these genes, *PvGATA17* was highly expressed in all tissues, followed by *PvGATA15/22. PvGATA19* had the least expression profile among the gene members, especially in roots. These findings reveal a unique and tissue-specific expression pattern of *PvGATA* TFs, which suggests their functional specialization in various tissues and developmental processes.

Based on previously generated RNA-seq data, we analyzed the expression profile of candidate *PvGATA* gene members in response to various abiotic and biotic stress conditions (Figure 6c, Table S4). Significant levels of positive and negative expression were observed among the *PvGATA* genes. Specifically, in both leaf and root tissues of common bean cultivars treated with saline water, *PvGATA* genes were preferentially expressed. Subfamily IV members showed the highest positive expression in leaf tissues, while subfamily II members were down-regulated. Notably, *PvGATA* genes displayed moderate expression in root tissues of resistant cultivars. In contrast, subfamily I members were up-regulated in lower hypocotyl tissues of salt-treated cultivars compared to the control. Similar expression patterns were observed for *PvGATA1/28*, whereas *PvGATA3/12/5* showed relatively lower expression levels. Subfamily II and III members exhibited relatively low expression, while subfamily IV had a moderately positive expression profile. Under dehydration stress in the drought-resistant cultivar Perola, subfamily IV members were predominantly up-regulated, with *PvGATA30/22/13/14/18* exhibiting the largest increase in up-regulation level. A similar expression pattern was observed for gene members of subfamily I and III, such as *PvGATA9/3* and *PvGATA7*, albeit to a lesser extent. In contrast, subfamily II gene members showed lower expression levels than all other genes. In drought-sensitive cultivars, a reverse expression pattern was observed across all genes, with *PvGATA30/7/3/18/29/22* being negatively expressed. This result suggests the potential importance of these genes in regulating resistance to drought stress in common bean plants. When exposed to fungal stress (*Sclerotinia sclerotiorum*), subfamily I members showed a significant increase in expression, with the highest expression value observed for *PvGATA10*. Subfamily III and IV members also exhibited increased expression levels, while the lowest expression levels were observed for *PvGATA8*, a member of subfamily II. *PvGATAs* displayed relatively low expression levels in response to cold stress across all subfamilies, with increased expression observed for *PvGATA17/22*. The most negatively expressed genes were *PvGATA10/14/28/3*, suggesting a potentially lesser role for *PvGATA* in modulating resistance to cold stress. In short, the expression patterns of *PvGATA* genes in different stress conditions suggest potential functional differences among the subfamilies.

To confirm the expression profiles of selected PvGATA genes, we performed expression analysis using qRT-PCR with RNA extracted from roots subjected to different abiotic and phytohormone treatments (Figure 7). Based on the expression levels of *PvGATAs* from previously analyzed RNA-Seq data (Figure 6c), we selected five *PvGATA* genes with the highest expression levels as candidate genes for further analysis. Under saline treatment, all analyzed genes were negatively expressed at 6 hours compared to the control. At 24hr, *PvGATA1, PvGATA10*, and *PvGATA28* were highly up-regulated, and *PvGATA25* was also positively expressed, albeit at a relatively lower level. At 48 hours, the expression levels were negative.

**Figure 7.**
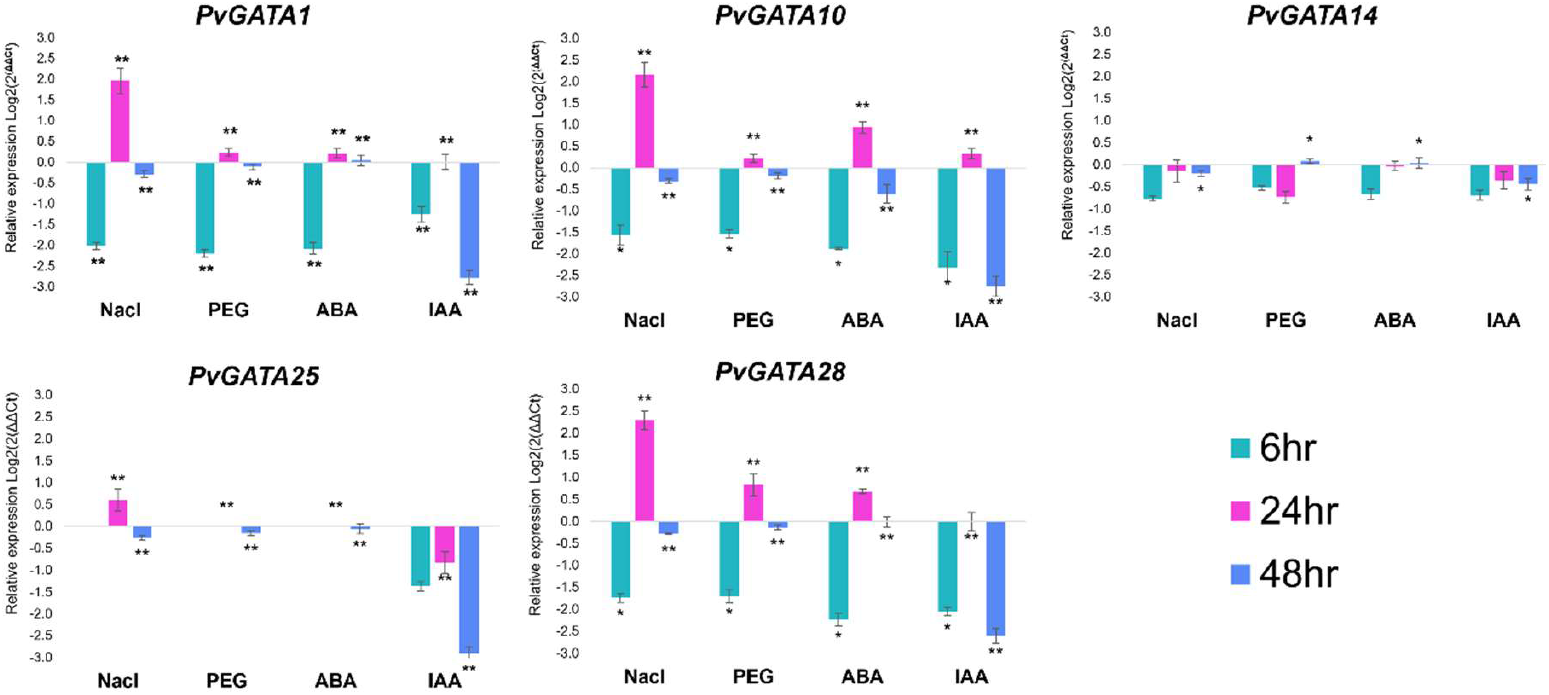
qRT-PCR expression profile analysis of five candidate PvGATA genes (PvGATA1, PvGATA10, PvGATA14, PvGATA25, and PvGATA28) under four different abiotic and phytohormone stress conditions, including, Salinity stress (NaCl), Drought stress (PEG), Abscasic acid (ABA), and Indole acetic acid (IAA). Samples were collected from root tissues at three time points after treatment, 6-hour, 12-hour, and after 48-hour. The experiments were performed independently with three replicates, and the error bars represent the standard deviation of three replicates. Asterisks indicate significant differences in transcript levels compared with the blank control. *P<0.05, **P<0.001.

Similarly, under drought treatment, all five genes were significantly down-regulated at 6 hours but peaked at 24 hours and showed low expression levels at 48 hours. Under phytohormone treatments with ABA (Abscisic acid) and IAA (Indole acetic acid), *PvGATAs* were generally negatively expressed, with a few exceptions. *PvGATA1, PvGATA10*, and *PvGATA28* were significantly up-regulated under ABA treatment compared to the control, and *PvGATA10* was up-regulated significantly under ABA and IAA treatment at 24 hours. Our results support findings from our previous RNA-seq analysis (Figure 6c) and suggest that *PvGATA1/10/25/28* may play crucial roles in regulating plant resistance against salt and drought stress at 24 hours. Additionally, these GATA genes may be involved in the phytohormone-mediated stress signaling pathways in the common bean plant.

## Discussion

The transcription factor *GATA* has been extensively studied in many plants for their regulating diverse crucial biological processes in plants by regulating the expression of genes responsible for the development, stress, and hormonal signaling (An et al., 2020; Manzoor et al., 2021; Reyes et al., 2004; Wang et al., 2023; Zhao et al., 2021). Common bean is a popular and commercially important crop, which has recently been affected by climate change. The functional and molecular mechanism of *GATA* TFs in regulating abiotic stress tolerance has not yet been studied in common bean plants. Hence, this study analyzed the functional identity of the *GATA* gene family in the *Phaseolus vulgaris* genome under different abiotic and phytohormonal stress.

In this study, we systematically analyzed the *P. vulgaris GATA* gene family. We identified 31 *GATA* TFs, a number close to that found in *Arabidopsis thaliana* (29), *Oryza sativa* (28) (Reyes et al., 2004), *Brachypodium distachyon* (27) (Guo et al., 2021), and *Capsicum tetragonum* (28) (Yu et al., 2021). More than those found in *Eucalyptus urophylla* (23) (Du et al., 2022) and *Prunus avium* (18) (Manzoor et al., 2021) and less than those found in *Triticum aestivum* (79) (Feng, Yu, Zeng, He, & Liu, 2022) and *Brassica napus* (96) (Zhu, Guo, Chen, Wu, & Jiang, 2020). Comparing *GATAs* of common bean to *Arabidopsis* and rice for collinearity analysis. We found that the collinearity between common bean and *Arabidopsis* was higher than in rice. This shows that number and function of *GATAs* are closely associated with species type. Among the 31 *PvGATAs*, 18 were duplicated, including six dispersed duplicated pairs, two whole genome duplicated pairs and one transposed duplicated pair. These results suggest the importance of gene duplication in contribution to *GATA* gene expansion and corresponds with previous findings in pepper, *Gossypium*, and apple (H. Chen et al., 2017; Yu et al., 2021; Z. Zhang et al., 2019). Furthermore, in previous reports phylogenetic analysis of *GATA* TFs identified seven subfamilies according to their conserved motif structure. In *PvGATAs* we grouped the genes into four subfamilies which lacks subfamilies V, VI, and VII which were previous reported in rice (Reyes et al., 2004), suggesting the conserved structure and function of *GATA* Tfs in plants. In our study, we identified eight, three, six and 13 members in each subfamily (subfamilies I. II, III, and IV) (Figure 3, Figure 2). A close relation in motif conserved domain and exon: intron structures was observed among same subfamily members suggesting their close functionality, but there have yet any reports on the functional differences among these subfamilies.

Most *GATA* Tfs in plants contain a single zinc finger domain in their protein sequence, such as those found in *Arabidopsis*, rice, wheat, and grape (Feng et al., 2022; Reyes et al., 2004; Z. Zhang et al., 2018). A single GATA domain was also present in all PvGATA protein sequences, however, members of subfamily II contained additional domains like CCT and tify domains. Previous studies also observed this and suggest a key role in diverse biological functions. For example, in regulating embryo and flower development, stress tolerance, and different phytohormone signaling (Behringer & Schwechheimer, 2015; Gupta, Nutan, Singla-Pareek, & Pareek, 2017; Richter, Behringer, Zourelidou, & Schwechheimer, 2013).

GATA factors were first predicted and identified in regulating light-associated genes and phytohormonal-regulated photomorphogenesis in *Arabidopsis* and *P. edulis* (Luo et al., 2010; H. Zhang et al., 2018). Hence, we predicted and analyzed the promoter cis-regulatory elements (CREs) in the 2000 bp upstream of PvGATAs. The results predicted a total of 799 CREs in all the PvGATAs. These CREs were categorized into three categories, namely growth and development regulatory elements, stress-responsive elements, and phytohormone-responsive elements. The growth and development regulatory elements had the largest share of CREs in the promoter region of PvGATA genes, with a total of 457 elements and 57.20% of total CREs. Within this category, CREs light responsiveness was heavily abundant, totaling 424 elements and 93% of the category’s total elements. This was similar to studies on cucumber and *Rosaceae sp* (Manzoor et al., 2021; K. Zhang et al., 2021). Other elements in this category included meristem expression elements, endosperm expression, circadian control, and seed-specific elements. The second category is stress-responsive CREs, with 151 (18.9%) CREs predicted in the promoter region of *PvGATAs*. This category includes anaerobic induction elements, wound-responsive elements, defense and stress responsiveness, drought inducibility elements, and low-temperature responsiveness. Anaerobic induction elements were most abundant within this category, indicating the vital role of *GATAs* in environmental stress resistance in common bean plants. The third category of CREs identified in the promoter region of *PvGATAs* is related to phytohormone response, with a total of 191 (23.9%) predicted CREs. The most common phytohormone-responsive element is the abscisic acid responsiveness element, followed by methyl jasmonate (*MeJA*)-responsive elements, salicylic acid-responsive elements, gibberellin-responsive elements, and auxin-responsive elements. Interestingly, subfamily II lacks CREs associated with gibberellin, auxin, and salicylic acid-responsive elements, suggesting the absence of these genes in the hormone regulation network. The identification of CREs in the promoter region of *PvGATAs* provides insight into the regulation of gene expression, particularly in growth and development, stress response, and phytohormone signaling. This information can be used to study gene expression and regulation, which can aid in crop improvement and stress tolerance. Understanding the functions of these regulatory elements in *PvGATA* genes can help identify molecular targets for crop improvement and breeding.

The GATA gene family encodes transcription factors that regulate gene expression by recognizing a specific consensus sequence NGATAY (N= T or A; Y= G or A (Jason A Lowry & William R Atchley, 2000). Nitrogen levels significantly impact plant growth and carbon uptake (Guerrieri et al., 2011), with low levels generally benefiting these processes, while high levels may decrease water use efficiency (Lu et al., 2018). In our study, we used PPI analyses to determine the roles of specific PvGATA proteins. PvGATA10, PvGATA12, PvGATA17, PvGATA28, and PvGATA29 play positive roles in nitrogen compound metabolic processes. Overexpression of these proteins could increase nitrogen usage, potentially counteracting the negative effects of high nitrogen levels in the soil. In addition to their roles in nitrogen metabolism, we also observed interactions between some PvGATA proteins and plant hormone signaling pathways. Plant hormone signaling is essential for responding to biotic and abiotic stressors, and our findings suggest that PvGATA proteins may play a role in these responses. For example, some studies have linked reduced cytokinin signaling to increased drought tolerance (Z. Liu, Yuan, Song, Yu, & Sundaresan, 2017; Nishiyama et al., 2013). In another study, the knockout of the gene that encodes the DELLA protein, a negative gibberellic acid signaling pathway regulator, has resulted in salt sensitivity (Achard et al., 2006). Conversely, overexpression of the gibberellic acid-insensitive-1 gene from *Arabidopsis* in Petunia has been linked to increased drought tolerance (Y. Zhang, Norris, Reid, & Jiang, 2021). Therefore, it is understood that reduced gibberellic acid signaling leads to increased drought tolerance and decreased salinity tolerance. Our study identified PvGATA10 and PvGATA28 as negative regulators of the gibberellic acid signaling pathway, suggesting that overexpression of one of these genes could lead to increased drought tolerance. This finding supports the potential use of these genes in developing drought-tolerant plants. Overall, our study provides valuable insights into the roles of specific PvGATA proteins in nitrogen metabolism and plant hormone signaling, offering potential avenues for future research on plant stress responses and crop improvement.

The expression profile of candidate *PvGATA* gene members in response to various abiotic and biotic stress conditions has been analyzed based on previously generated RNA-seq data. The study revealed that *PvGATA* genes exhibited a significant level of positive and negative expression among the subfamilies in response to different stress conditions. The subfamily IV members showed the highest positive expression in leaf tissues. In contrast, subfamily II members were downregulated in both leaf and root tissues of common bean cultivars treated with saline water. Subfamily I members were up-regulated in lower hypocotyl tissues of salt-treated cultivars compared to the control. *PvGATA* genes displayed moderate expression in root tissues of resistant cultivars, and subfamilies III and IV had a moderately positive expression profile under dehydration stress in the drought-resistant cultivar Perola. In contrast, subfamily II gene members showed lower expression levels than all other genes in drought-resistant and drought-sensitive cultivars. This suggests that subfamily IV members are potentially important in regulating resistance to drought stress in common bean plants. Under fungal stress, subfamily I members showed a significant increase in expression, with *PvGATA10* showing the highest expression value. Subfamily III and IV members also exhibited increased expression levels in *response* to fungal stress. The study confirmed the expression profiles of selected PvGATA genes using qRT-PCR with RNA extracted from roots subjected to different abiotic and phytohormone treatments. The results were consistent with the previously analyzed RNA-Seq data, suggesting that *PvGATA1/10/25/28* may be crucial in regulating plant resistance against salt and drought stress at 24 h after treatment. Additionally, these *GATA* genes may be involved in the phytohormone-mediated stress signaling pathways in the common bean plant.

Overall, the study suggests potential functional differences among the subfamilies of *PvGATA* genes in different stress conditions. Identifying specific *PvGATA* genes that are up or down-regulated in response to stress may provide a foundation for breeding common bean plants with improved stress tolerance. Additionally, the study highlights the potential importance of the *GATA* gene family in regulating stress responses in common bean plants and provides insights into the molecular mechanisms underlying stress tolerance in plants.

## Conclusion

In conclusion, this study provides insights into the functional identity of the GATA gene family in common bean plants. 31 GATA transcription factors were identified in *P. vulgaris*, this number was close to that found in Arabidopsis and rice. Gene duplication is also important in the expansion of the *GATA* gene family in common bean, and our phylogenetic analysis identified four subfamilies of *PvGATAs*. Members of subfamily II contained additional domains, such as CCT and tify domains, which have been observed to play a key role in diverse biological functions in other plants. Our analysis of promoter cis-regulatory elements predicted 799 elements in all the *PvGATAs*. Five genes were selected for qRT-PCR expression analysis, results indicated that PvGATAs may play role in initial expression under abiotic stress. This study provides a basis for further functional studies on *PvGATAs* in regulating abiotic stress tolerance and growth and development in common bean plants.

